# Analyses of transposable elements in arbuscular mycorrhizal fungi support evolutionary parallels with filamentous plant pathogens

**DOI:** 10.1101/2024.11.04.621924

**Authors:** Jordana I. N. Oliveira, Catrina Lane, Ken Mugambi, Gokalp Yildirir, Ariane M. Nicol, Vasilis Kokkoris, Claudia Banchini, Kasia Dadej, Jeremy Dettman, Franck Stefani, Nicolas Corradi

## Abstract

Transposable elements (TEs) are repetitive DNA sequences that excise or create copies that are inserted elsewhere in the genome. Their expansion shapes genome variability and evolution by impacting gene expression and rearrangement rates. Arbuscular mycorrhizal fungi (AMF) are beneficial plant symbionts with large, TE-rich genomes, and recent findings showed these elements vary significantly in abundance, evolution, and regulation among model AMF strains. Here, we aimed to obtain a more comprehensive understanding of TE function and evolution in AMF by investigating assembled genomes from representatives of all known families. We uncovered multiple, family-specific bursts of insertions in different species, indicating variable past and ongoing TE activity contributing to the diversification of AMF lineages. We also found that TEs are preferentially located within and around candidate effectors/secreted proteins, as well as in proximity to promoters. Altogether, these findings support the role of TEs in promoting the diversity in proteins involved in molecular dialogues with hosts and, more generally, in driving gene regulation. The mechanisms of TEs evolution we observed in these prominent plant symbionts bear striking similarities to those of many filamentous plant pathogens.

## Introduction

Arbuscular Mycorrhizal Fungi (AMF) are important plant symbionts in the sub-phylum Glomeromycotina[1]. These organisms support their hosts by transferring nutrients from the soil in exchange for carbohydrates and lipids, essential to their metabolism[2–4]. Fossil and phylogenomic evidence indicates that this successful partnership occurred over 400 million years ago when plants colonized land[2,5]. AMF hyphae and spores carry hundreds to thousands of nuclei[6], and these organisms have long been thought to be asexual due to the absence of formal sexual reproduction or structures[7]. However, this assumption has been challenged by the discovery of genes involved in sexual reproduction[8–10] and, more recently, by the discovery of homo-heterokaryotic life stages[11,12].

Genome size in AMF varies from about 50 to over 800 Megabases (Mb)[13], and correlates with genome composition, with larger genomes carrying significantly more repeated sequences, particularly transposable elements (TEs)[14]. TEs are repetitive DNA sequences that self-synthesize and insert copies of themselves within a genome, shuffling genetic information and creating functional novelties by inserting nearby promoters to impact gene expression[15] or in coding regions to alter protein sequences[16,17]. These elements are divided into two major classes based on their mobilization mechanisms: Class I (retrotransposons) use a “copy-and-paste” mechanism via an RNA intermediate to create new insertions, whereas Class II (DNA transposons) promote insertions by “cutting and pasting” DNA sequences[18]. TE expansions are controlled by DNA methylation and RNAi to maintain genome integrity. Still, external pressure, such as stress and variation of abiotic factors, can repress these silencing mechanisms and thus activate TE expansion[19]. Over time, TEs can also be domesticated to create new functions, a process known as molecular cooption, leading to increased protein variability that promotes adaptation to environmental changes[20].

In AMF, TE diversity has been detailed in a few species, mainly in the Gigasporaceae and Glomeraceae families[21,22]. More recently, their potential functional role has been uncovered in the model species *Rhizophagus irregularis*, where TE expression differs across life stages and is significantly up-regulated upon plant colonization[23,24]. Notably, TE families are differentially regulated depending on plant host identity, suggesting that these elements play an active, yet unknown, host-specific role during colonization[23].

Significant differences in TE abundance, genome location, and evolution exist between *R. irregularis* strains and species[13,23]. For instance, in the *R. irregularis* strain DAOM197198, the mobilization and retention of TEs can differ between related strains, and different species carry unique TE expansions, such as the long interspersed nuclear elements (LINEs) retroelements in *Gigaspora margarita*, which have been hypothesized to drive genome expansion in this genus[21,25]. TE insertion/retention also varies within AMF genomes. For example, in *R. irregularis*, the genome separates into two functionally distinct regions referred to as A- and B-compartments[12,26]. Most of the conserved genes are in the A-compartment, while the B-compartment is more repeat-dense and carries most genes potentially involved in the dialogue with the plant – i.e. secreted proteins (SP) and candidate effectors. Young TEs accumulate at higher rates in the latter but are eliminated early from the genome in the A-compartment[23].

It has been hypothesized that the abundance of TEs within the B-compartment in AMF mirrors what is observed in the genomes of some filamentous fungal and oomycete plant pathogens[24,26,27]. Specifically, these genomes can also separate into slowly evolving regions carrying conserved genes (analogous to the A-compartment), and others that evolve rapidly due to significantly higher TE and effector density (analogous to the B-compartment), leading to enhanced diversification of proteins involved in virulence[28,29]. As in some plant pathogens, B-compartment effectors are up-regulated in AMF upon colonization[26], and a significant positive correlation in the expression of TEs and B-compartment genes has been identified in the AMF *R. irregularis*[23].

Despite apparent similarities in the mode of TE evolution and regulation between AMF and distant plant pathogens, the mechanisms behind these have not been fully investigated. Furthermore, it is unknown whether the similarities observed between model AMF and plant pathogens are conserved among all Glomeromycotina lineages, as a detailed analysis of TEs in most known families has not yet been performed[23]. Here, we aim to improve our understanding of the diversity and evolution of Glomeromycotina TEs in comparison to pathogenic relatives by analyzing 24 AMF species with available genome assemblies, encompassing all known orders of this prominent symbiont group[30,31].

## Results

### Transposable elements diversity in Glomeromycotina

Transposable elements were annotated in 24 genome assemblies, including 3 Paraglomerales, 3 Archaeosporales, 5 Glomerales, 11 Diversisporales, and 2 Entrophosporales (**Table 1**). Only *Rhizophagus irregularis* have chromosome-level assemblies [26,32]. The quality of the other assemblies varies from a N50 of 1,501 to 1,306,809 in *Innospora majewskii* and *Rhizophagus clarus*, respectively (Table S1). To aid visualization of TE evolution, a phylogenetic tree was generated using OrthoFinder [33] that fully agrees with the most recent phylogenomic analyses of AMF (**Figure S1**) [34].

**Table 1.** Comparison between automated and customized TE annotation.

| Group | Species | $\bar{X}\%$ masked TE before | $\bar{X}\%$ unknown TE before | $\bar{X}\%$ known TE before | $\bar{X}\%$ known TE after |
| --- | --- | --- | --- | --- | --- |
| Paraglomerales | 3 | 22.3 ± 2.2 | 18.4 ± 2.1 | 3.9 ± 2.15 | 8.6 ± 0.32 |
| Archaeosporales | 3 | 62.5 ± 6.66 | 18.8 ± 6.75 | 43.7 ± 5.47 | 49.2 ± 9.99 |
| Glomerales | 5 | 45.8 ± 3.32 | 30.5 ± 1.55 | 15.3 ± 1.3 | 30.5 ± 2.85 |
| Diversisporales | 11 | 68.5 ± 2.62 | 45.3 ± 2.59 | 23.2 ± 0.98 | 43.9 ± 2.11 |
| Entrophosporales | 2 | 34.3 ± 0.3 | 25.8 ± 0.45 | 8.6 ± 0.38 | 14.1 ± 0.82 |

The mean percentage of the genome ( ± SE) covered by known or unknown TEs in each order was calculated for each of the two annotations. In the first annotation, the masked portion represents known and unknown TE sequences. There is an increase in known sequences for all orders after the curation. The highest and lowest mean percentages of known sequences are shown in green and red, respectively.

Using the automated method RepeatModeler to generate the repeat libraries, between 18% to 45% of genomes were covered by unknown TEs. This high number of unclassified repeats is mainly due to the lack of information on TEs from fungi in the main databases used as reference[35]. Building a customized curation over the automated repeat library, significantly improved the annotation of transposable elements in all genomes (p <0.01), except for *Geosiphon pyriformis* and *Acaulospora colombiana*, where most TEs were already derived from known sequences. With this curated dataset, the proportion of TEs within AMF genomes now ranges from a minimum of 6.82% in *Innospora majewskii* to a maximum of 58.98% in *Ambispora leptoticha* (**Figure 1**). AMF genomes also contain several expanded gene families, such as Kinases, Kelch, Sel1, TPR, and BTB, which represent on average 3.7% of the AMF curated datasets, with *R. irregularis* containing the highest number of repeated genes at 8%[24,36] (**Figure 1a**).

**Figure 1.**
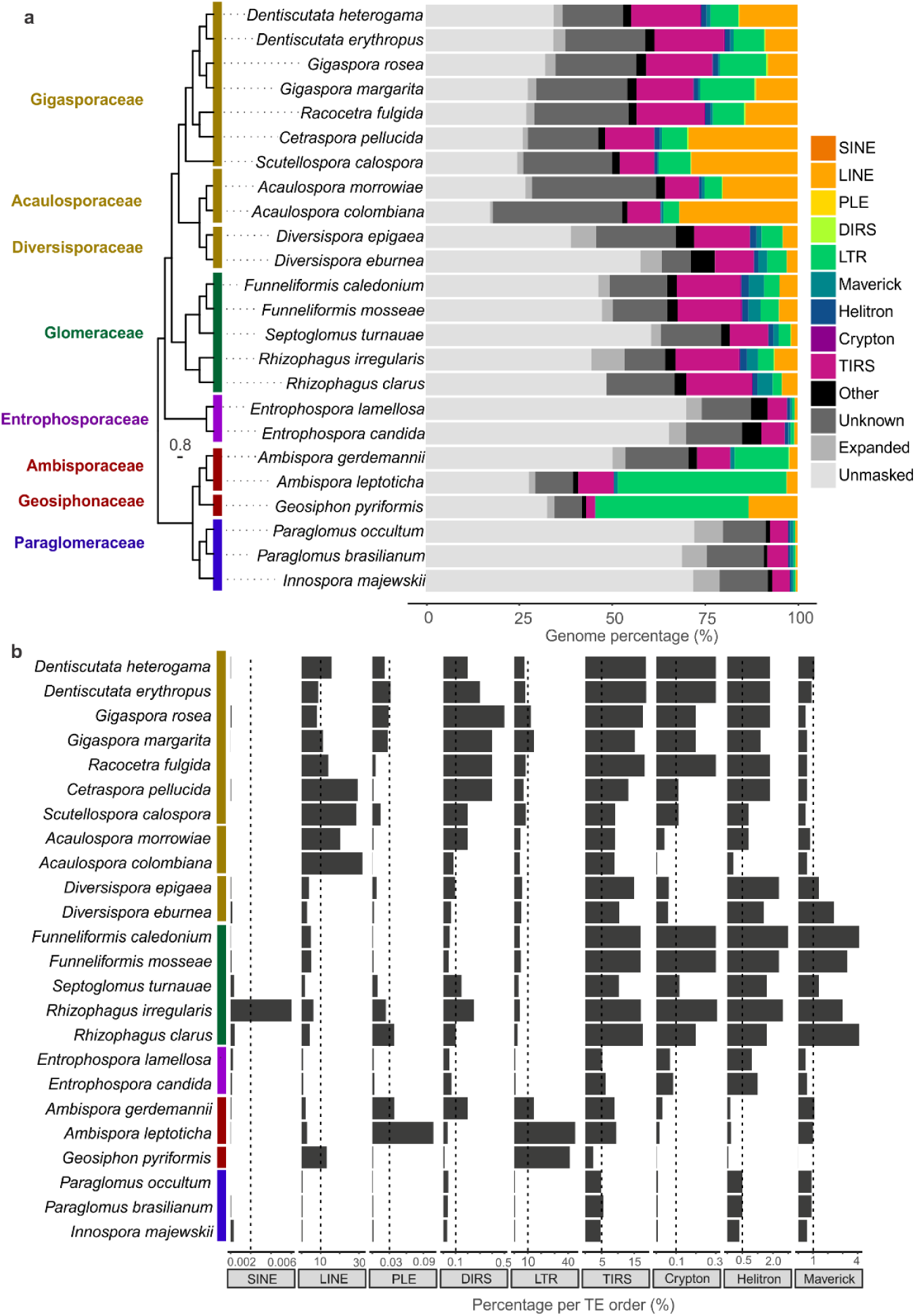
Genome composition in Glomeromycotina species. **a)** Relative distribution of different types of sequences in AMF species. The phylogenetic tree is based on ortholog genes, and the bar indicates the level of nucleotide substitution, representing the distances between the taxa. Bootstrap support can be found in Figure S1. The tree is consistent with the most AMF phylogeny built by Rosling et al. 2024. The colors on the genome composition represent different TE orders. In black, are other repeats, such as satellite and low complexity; in dark grey are the expanded gene families. The “unknown” buffer represents the remained portion of masked repeats from the first annotation (without curation). The “unmasked” region represents the genome portion other than repeats. **b)** Relative content per order of transposable element in relation to the entire genome. The *x-axis* shows the genome percentage covered by each TE order.

In general, species within the same order share very similar TE diversity. Paraglomerales species always have the lowest content of TEs, ranging from 6.8% to 8.1%. The Archeosporales carry long-terminal repeats (LTRs) expansions especially *G. pyriformis* with 41.2% of the genome covered by LTR sequences. In the Glomerales, DNA transposons dominate TE abundance and carry unique expansions of Maverick elements. The expansion of LINE elements found in *Gigaspora* species, previously hypothesized to be the cause of genome expansion in these genera[21,22], is common to all Diversisporales species (i.e. Acaulosporaceae, Diversisporaceae, and Gigasporaceae families).

### A core of highly expanded TEs in Glomeromycotina

Of the 6,772 non-redundant consensus sequences curated in this study, 1,295 (19.12%) are shared among all species (“core” families) (**Figure 2a**), an indication that the 4,477 sequences either originated and diversified after species differentiation, or were lost secondarily in different lineages. The AMF “core” families represent on average 50% of the TEs copies in AMF genomes (**Figure 2b**). Among the core sequences, LTR/Bel has two families that are present in all genomes analyzed. In contrast, LTR/Copia, DNA/Academ, DNA/Dada, Crypton, and SINE are the superfamilies with a higher number of sequences that are “non-core” families.

**Figure 2.**
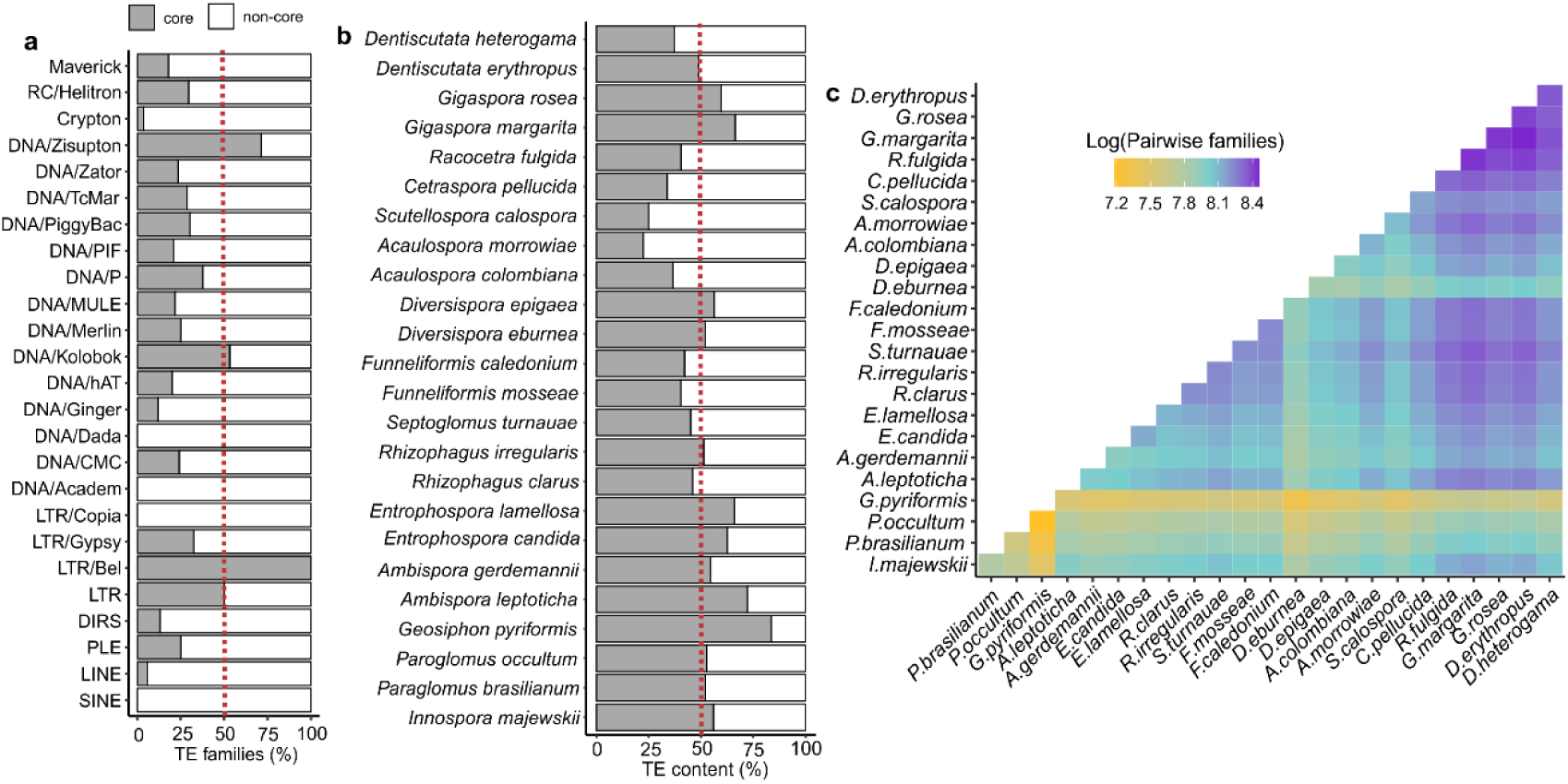
TEs families shared across Glomeromycotina species. **a)** Percentages in relation to the total number of each superfamily that are present in all species (core) and in blank are the superfamily that are not present in all species (specific). **b)** Relation of core and specific families in the TE annotation. **c)** Pairwise comparison of shared families. The number of shared families between two species is expressed in log scale to minimize the bias of certain species having more families.

Overall, species from Glomerales and Diversisporales have more expansions of clade-specific sequences and carry a greater number of non-core families compared to the other AMF groups. Pairwise comparisons showed that the non-core families are more likely to be shared between closely related species, and could represent clade-specific TE invasions and expansions (**Figure 2c**).

### Transposable elements expansions and degeneration across AMF species

To determine how TEs are retained and mobilized in AMF, we constructed their repeat landscapes based on the Kimura substitution calculation. Lower Kimura substitution rates indicate recent insertion events, with sequences showing a low level of divergence from the consensus, whereas higher Kimura substitution rates uncover older TE insertions, meaning a high-level divergence between the copies and the consensus.

Clear patterns of conservation and distinctions in TE mobilization and retention exist among analyzed families (**Figure 3**). Specifically, the Paraglomeraceae and Entrophosporaceae have a flat distribution of TEs (**Figure 3a**), suggesting that minimal TE bursts have occurred in these two distinct lineages. In contrast, other families show specific waves of TE expansion and degeneration leading to genome diversification. For example, Geosiphonaceae and Ambisporaceae underwent similar and striking expansions of LTRs, while LINEs have similarly invaded Acaulosporaceae and Gigasporaceae. Of all the AMF families analyzed, Glomeraceae and Diversisporaceae were the only ones to show obvious recent expansions (peaks between 1 to 5 sequence divergence) (**Figure 3e, f**), suggesting ongoing TE activity in their genomes.

**Figure 3.**
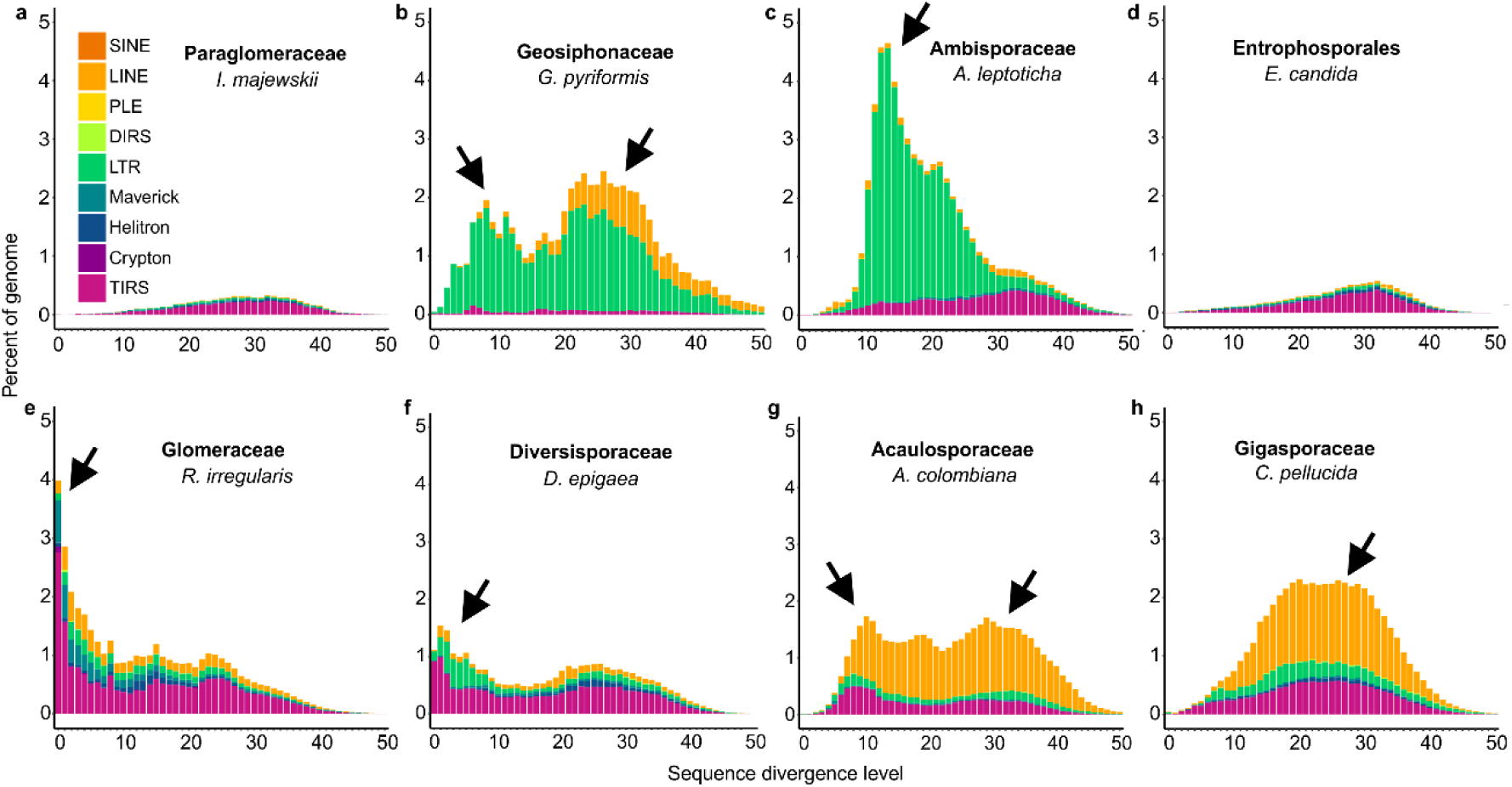
Transposable elements landscape across AMF species. Each histogram represents the genome percentage (y-axis) of copies with different level of divergence (x-axis). The arrows highlight the specific past events of TE that might contributed to the diversification of the genome.

### TEs are preferentially located upstream of genes in Glomeromycotina

In filamentous plant pathogens, TEs located within 1000 bp upstream to their transcript start site (-1Kb TSS), - i.e. in obvious vicinity to promoters and/or regulatory sequences that influence gene expression - are co-express with the genes they co-locate with [37,38]. In the model AMF *R. irregularis*, genes within the -1Kb TSS also co-express with the closest TEs [23].

By investigating available datasets, we confirm that, in all AMF orders, the closest TE to a gene locates within the 1Kb-TSS (average -872 nucleotides upstream of the TSS, (SD = 399) (**Figure 4a**). In all cases, TE insertions are significantly more common within -1Kb TSS regions compared to both non-coding regions (UTRs and introns, average of 38% *vs* 20%; p-value <0.001) and exons (average 38% *vs* 30%; p-value <0.001) (**Figure 4b**). For the -1Kb TSS insertions, notable differences exist among families (SD 17%). For example, 20% of genes in Paraglomerales 60% of TEs are within the TSS compared to up to 60% in Glomerales and Diversisporales. Overall, *Paraglomus* spp. and *A. colombiana* have fewer genes with TE sequences within CDS/exons at 13% and 9% respectively, whereas *D. epigea* and *G. pyriformis* have a higher proportion with 53% and 61% of TEs. *G. pyriformis* is also characterized by a much higher number of genes with TEs insertions within exons (**Figure 4b**). In Glomeromycotina, LINE, LTR/Gypsy, DNA/CMC, DNA/hAT, and DNA/MULE elements are the TE families commonly found either near or within genes (**Figure S2**).

**Figure 4.**
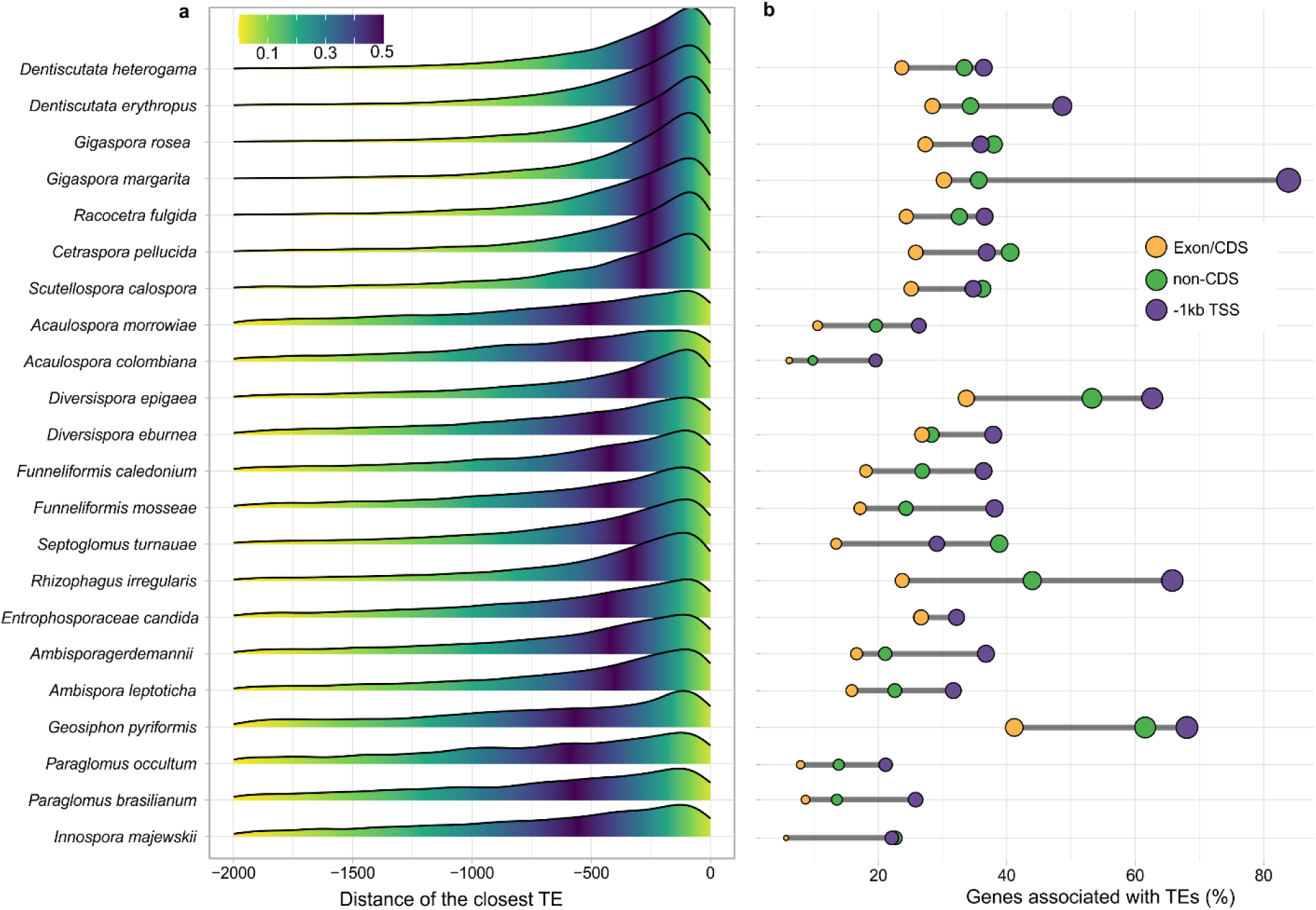
Transposable elements association within genes. **a)** Closest TE association in the upstream region of coding genes. The x-axis represents the base-pair distance from the transcription start site (TSS). The y-axis shows the density of genes with the closest TE at a given position (x-axis). The gradient color indicates the tail probability, highlighting regions with significant TE proximity. **b)** Percentages of genes harboring TE insertions upstream to the transcription start site (TSS), in UTRs and introns (non-CDS) and exon/CDS regions.

### Effectors locate within TE-rich regions in Glomeromycotina

The effector genes of several filamentous plant pathogens are often associated with TEs within so-called “adaptive regions” that promote effector variability and diversification[39]. These genes are important to establish communication with the host and start the infection process.

With the exception of *P. occultum*, *G. pyriformis,* and *A. marrowiae*, the density of TE insertions, defined by sequence overlaps with coding sequence and co-location within 1Kb up or downstream genes, is significantly higher (p-value <0.05) within candidate effectors compared to other protein-coding genes **(Figure 5, Table S4**). Effectors are predominantly found in TE-rich regions, while non-secreted genes are more common in TE-poor areas, with secreted proteins showing an intermediate pattern. In most species, effectors exhibit a right-skewed TE frequency distribution, supporting genome compartmentalization where dynamic, TE-dense regions harbor SPs and effector genes. This pattern aligns with the “two-speed genome” model observed in plant pathogens [35,37,38,40], and the A/B genome compartments reported in the model AMF *R. irregularis* [12,23,26].

**Figure 5.**
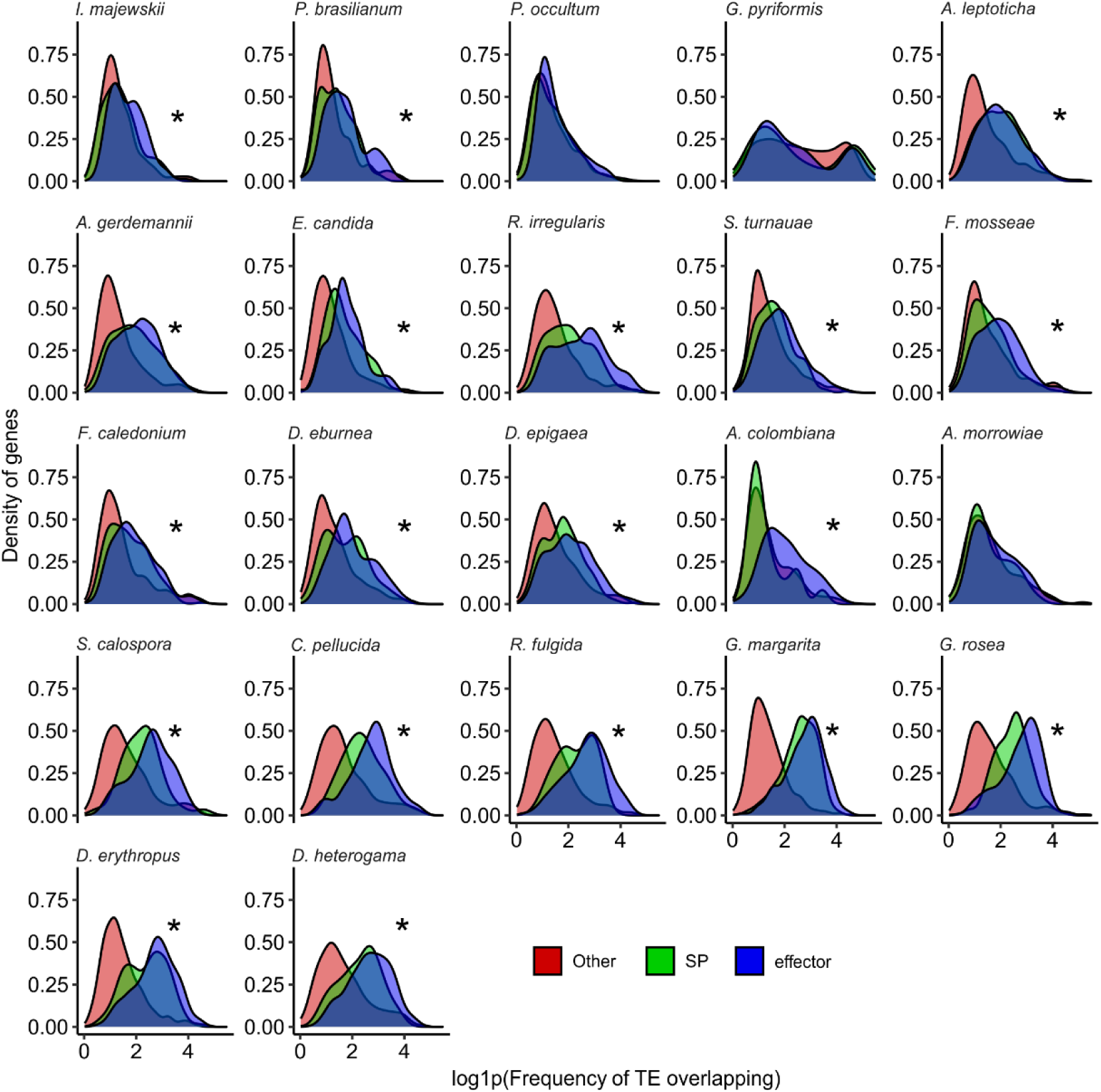
TE overlap frequency within and around Genes. The x-axis represents the log-transformed frequency of TE insertions overlapping gene regions, while the y-axis shows the density of genes with a given TE overlap. Effectors (blue) are compared to secreted proteins (SP, green) and other protein-coding genes (red). Asterisks (*) indicate significant differences (t-test p < 0.05) between effectors and other genes. For additional statistical comparisons, see Table S4.

It is noteworthy that the degree of fragmentation of the genome assembly does not affect the general observations; in almost all cases, TEs are more likely to be found near or within promoter regions (-1Kb) and in association with secreted proteins and effectors. Even species with the lowest BUSCO scores (<50%), such as *Scutellospora calospora, Dentiscutata heterogama, Entrophospora lamelosa,* and *Innospora majewskii*, presented the observed trends are like those identified in more complete assemblies (**Table S1**).

## Discussion

### A curated view of TE diversity and evolution in the Glomeromycotina

The current work provides a curated catalogue of TEs for all Glomeromycotina with available genome assemblies. Although the curation improved the TE annotations performed with tools based on sequence similarity[41,42], most of the families we analyzed, except *R. irregularis*, still have a proportion of putative unknown repeats. This is likely because *R. irregularis* is the only AMF species with chromosome-level assemblies, while others have fragmented assemblies or collapsed repeat regions[30], which are known to limit the detection of TEs annotation[35]. Hopefully, more chromosome-level assemblies will become available in the future, allowing the identification of species-specific TEs that may have been overlooked in our screening.

In general, AMF and other fungi carry expansions of retrotransposons such as LINEs and LTRs, however, some species deviate from this trend[21,35]. For example, insertions from DNA transposons such as TIRS, Crypton, Helitron, and Maverick elements are abundant in Glomerales. The patterns of TE evolution we observed also confirm that, together with the number of genes[13], TEs can vary dramatically in presence/absence and relative abundance between AMF relatives and within TE families. For example, the virus-like element Maverick is expanded exclusively in Glomerales. This element has only been reported in the plant pathogen *Phakosora pachyrhizi*, in the AMF *Rhizophagus (Glomus) intraradices*[43], and in the chytrid *Batrachochytrium dendrobatidis* [44], and it is thus intriguing to speculate that it may have originated via horizontal gene transfer in the Glomerales. Another, perhaps less realistic scenario, is that this TE family was lost in all but one of known AMF orders.

In general, our work supports the view that obligate biotrophs carry more transposable elements than non-obligate fungi, linking the pattern of TE insertions to their lifestyle[35,45]. Within this context, a similar trend was observed in the cyanobacteria endosymbiont *G. pyriformis*, which showed a greater expansion of mobile genetic elements compared to any free-living relatives[46].

### TE diversity and distribution often reflect family evolutionary history

In many cases, TE diversity and distributions were very similar between members of the same AMF family. This supports events of TE insertions and retention that occurred in the most recent common ancestor (MRCA) of these lineages. For example, all members of the Archaeosporales that we analyzed showed similar bursts of LTRs and differed only in how each of these TEs was retained over time. Similarly, members of the sister orders Glomerales and Diversisporales underwent strikingly similar bursts of LINEs and DNA/TIRs, again suggesting that these were inserted before these lineages diverged.

Remarkably, the evolution of these elements can sometimes go beyond phylogenetic relationships. For example, members of phylogenetically distinct families, Paraglomeraceae and Entrophosporaceae, can have virtually identical TE distributions. This similarity may reflect independent and adaptive TE retentions due to similarities in (epi) genomic architecture, such as methylation, or ecological traits[35].

### Evidence of A/B compartments across all Glomeromycotina

The chromosomes of the model species *R. irregularis* are separated into two main regions, called A/B compartments[12,26], and our analysis strongly indicates that this organization is shared by most Glomeromycotina. In particular, TE density is always significantly higher near secreted proteins and candidate effectors in AMF genomes; a characteristic that is highly reminiscent of the B-compartment and that shares further highlights striking similarities in the genome biology of AMF and many filamentous fungal plant pathogens[24,29].

Specifically, these independent symbiotic and pathogenic lineages both carry genes involved in dialogue with the host (effectors) within fast-evolving and repeat-rich regions of the genome. As in fungal pathogens, the association between effectors and TE in AMF is likely to diversify these molecules, allowing for improved adaptation and symbiosis with a wider range of hosts. Notably, this organization can also lead to a “devil’s bargain” situation, as TE insertions near or within effectors could degenerate these genes by increasing the chances of deleterious mutations for the symbiosis[39].

Future studies based on analyses chromosome-level assemblies and chromatin-capture (Hi-C) information from representatives of the orders Diversisporales, Archeasporales, and Paraglomerales will hopefully provide a more detailed view of the shared and lineage-specific regulatory processes affected by each compartment in Glomeromycotina.

### Are TE bursts linked with AMF species diversification and adaptation?

Bursts of TEs promote genomic rebuilding, and this mechanism is often associated with higher speciation rates[47]. In this context, it is intriguing to speculate that the reduced TE load of Paraglomerales, together with its flat landscape distribution (indicating low TE activity and highly stable genome) is linked with a limited species richness within this family[48]. Indeed, to date, Paraglomerales has fewer species formally recognized compared to other Glomeromycotina, including other early diverging lineages like Archaeosporales – e.g. 11 Paraglomerales and 19 Archaeosporales[48]. In contrast, more divergent AMF orders such as Glomerales and Diversisporales featured high levels of TE bursts within their genomes and are significantly more species-rich, with 268 and 70 species, respectively[48].

The lower TE abundance of Paraglomerales also supports the hypothesis that fewer gene/TE associations with reduced mycorrhizal capabilities[14] recently reported for this order compared to AMF relatives[49,50]. Here, lower TE loads may reduce the chances of generating genomic diversity necessary for adapting to environmental change[27,51], especially if mating is rare or absent[10,24]. In support of this, most other AMF groups with high genomic diversity and adaptability[13], such as Glomeraceae, have experienced notable TE bursts in the past and carry evidence of recent and ongoing TE activity.

### Indirect evidence of gene regulation through TE in Glomeromycotina

Our analyses also revealed that TEs are preferentially locate within promoter regulatory regions in these symbionts. TE insertions near genes can impact gene expression by inducing methylation or altering transcription factor binding sites[52], and even promoting antifungal resistance as observed in a plant pathogen[53]. In the yeast *Saccharomyces cerevisiae*, LTR elements are preferentially inserted 700 bp upstream to coding genes and show synergistic expression during heat-stress induction[54].

In *R. irregularis*, co-expression has been observed between SP and upstream TEs in the B-compartment, with TEs showing higher expression during symbiosis[23] than in spores[24]. In addition, TEs showed strong methylation signals in germinating spores[26], which coincides with high expression of an RNAi-related gene[23]. This suggests that TEs may undergo demethylation during symbiosis, leading to increased expression of both TEs and symbiosis-related genes, such as secreted proteins and effectors. Again, this phenomenon has been observed in plant pathogens, where genes with TE insertions within -1Kb are highly expressed during early infection, revealing a synchrony between TEs and infection-related genes[37,54].

Remarkably, our analyses revealed that regions containing effector genes have significantly more TE insertions compared to other coding sequences. This phenomenon has also been observed in plant pathogens, where TE accumulation drives effector diversification, thereby increasing fungal virulence[28,39]. On the other hand, the plant evolves its immune system to combat infections, and again, TE-driven modifications serve as a source for the fungus to overcome host defenses, in an ongoing coevolutionary process[55]. Similarly, in AMF, diversified effectors could lead to broader adaptation to different hosts and environments, as these cosmopolitan symbionts can associate with many different hosts[56]. Given the high expression of TEs during symbiosis and frequent association with SP, it would be interesting to investigate changes in methylation to understand how these sequences become more actively transcribed upon contact with the host. Additionally, it would be valuable to examine which factors, whether biotic or abiotic stressors, can demethylate TEs in spores and reactivate TE mobilization as a source of variability.

As our study uncovered the presence of numerous species and order-specific TE families, it will be important that future analyses aim to determine their relative function and regulation among various conditions and plant hosts. This will be a crucial step to obtain a complete understanding of the evolution and role of TEs in these widespread root symbionts. Within this context, Maverick and LINE elements, which present specific expansions in Glomerales and Diversisporales, are good candidates for targeted investigation.

## Material and Methods

### Genome assemblies

We downloaded 21 genome assemblies, protein, and gene annotation files from all Glomeromycotina orders from NCBI and provided 3 new genome assemblies **Table S1**. Despite other AMF species having genomes deposited in the databases, we opted to keep only the assemblies with protein and gene annotation files which were used in the further analysis.

### Hidden-Markov Models (HMM) of transposition domains

We constructed HMM profiles for several transposition domains described in **Table S2**. Sequences for each domain were aligned separately using mafft[57], then converted to Stockholm format using esl-format and submitted to hmmbuild[58] to generate the HMM profiles. The profiles were concatenated and prepared for hmmscan using hmmpress. All models are available on https://github.com/jordana-olive/Transposable-Elements-in-AM-Fungi-Evolutionary-Parallels-with-Fungal-Plant-Pathogens.

### Repeat library generation

Each genome was submitted to RepeatModeler2.0[41] with the -LTRStruct option to generate the repeat sequence files. All the repeat libraries were concatenated into a single file from which sequences shorter than 1000 nucleotides were removed using seqkit [59]. The most representative sequence (consensus) for each putative TE family was prospected by clustering the repeat library using cd-hit-est -aS 0.8 -c 0.8 [60], following the 80-80-80 rule. This rule establishes that all sequences with at least 80% similarity along 80% of the total length should belong to the same TE family [61]. The final file containing the nonredundant TE-families was named “non-curated library”.

### Transposable elements curation and annotation

We predicted open reading frames (ORFs) in the “non-curated library” using getorf[62], selecting translated ORFs of at least 300 nucleotides. We submitted the ORFs from the “non-curated library” to hmmsearch [58] to retrieve the sequences containing the transposition domains using the hmm models. We selected the best match of each sequence using HmmPY.py[63] (e-value 1e-17, -c 0.35, -h 50). Sequences carrying transposition domains were renamed with the corresponding family domain. Curation generated the “final library” to be used for transposable element annotation. To compare the impact of the curation on the TE resolution, the annotation was performed twice in each genome using RepeatMasker2.0.4 -s -a modes[42] using the non-curated library and the final library. The alignment files generated by the annotation process were used to calculate the degree of divergence based on Kimura values using the scripts calcDivergenceFromAlign.pl and createRepeatLandscape.pl provided by the RepeatMasker tool.

### Orthologues genes and AMF phylogenetic tree

The protein sequences of the studied genomes were submitted to OrthoFinder[33] to construct a phylogenetic tree based on orthologous genes, and iTOL was used to visualize the branch length and bootstrap support (**Figure S1**)[64]. The tree was rooted with species from Paraglomerales and Archeosporales and visualized using the ggtree R package in combination with the TE annotation data.

### TE association with genes

We compared the intersection between TEs and genes using bedtools intersect[65] in gff from Genebank and our TE annotation. To maximize the results, we rely on the species with available gene annotation and protein sequences, which allow us to confirm the gene structure by comparing coding sequences. We also split the annotations into three files: 1000 base pairs (bp) upstream of the gene’s transcription target site, exons and other non-coding regions (5’ & 3’UTR and introns). For instance, using bedtools subtract, introns and UTRs were prospected by subtracting the coding region information following the protein sequences coordinates[65]. We considered a valid intersection when the TE overlap was at least 50 nucleotides long and not identical to the size of TE and the gene simultaneously.

### Prediction of secreted proteins and effectors

Secreted proteins were predicted in all species using SignalP-6.0[66]. From the predicted secreted proteins, we used the EffectorP-3.0[67] to annotate the candidate effectors.

## Acknowledgments

Our research is funded by the Discovery program of the Natural Sciences and Engineering Research Council (RGPIN2020-05643) and a Discovery Accelerator Supplements Program (RGPAS-2020-00033). NC is a Research Chair in Microbial Genomics at the University of Ottawa. JINO is funded by the Mitacs Accelerate Program (IT16902) and a Discovery Accelerator Supplements Program (RGPAS-2020-00033).

## Author’s contribution

Experiment design, contextualization and writing (JINO and NC). Bioinformatic analysis (JINO, CL, KM, GY). Repeat annotation and repeat landscapes (CL). Biological material, sequencing and analyses for *I. majewskii, S. turnauae* and *E. lamellosa* (CB, VK, FS, KD, JD). Data curation and downloading (AMN, GY). Github organization (KM). Supervision (NC). All authors read and approved the final version of the manuscript.

## Notes

https://github.com/jordana-olive/Transposable-Elements-in-AM-Fungi-Evolutionary-Parallels-with-Fungal-Plant-Pathogens

https://doi.org/10.5281/zenodo.13984052

### Competing Interest Statement

The authors have declared no competing interest.

### Summary of Updates

Addition of figures and analyses further supporting strong association between TEs and specific locations/genes in the genome

