## Supplemental figures for "Analyses of transposable elements in arbuscular mycorrhizal fungi support evolutionary parallels with filamentous plant pathogens"

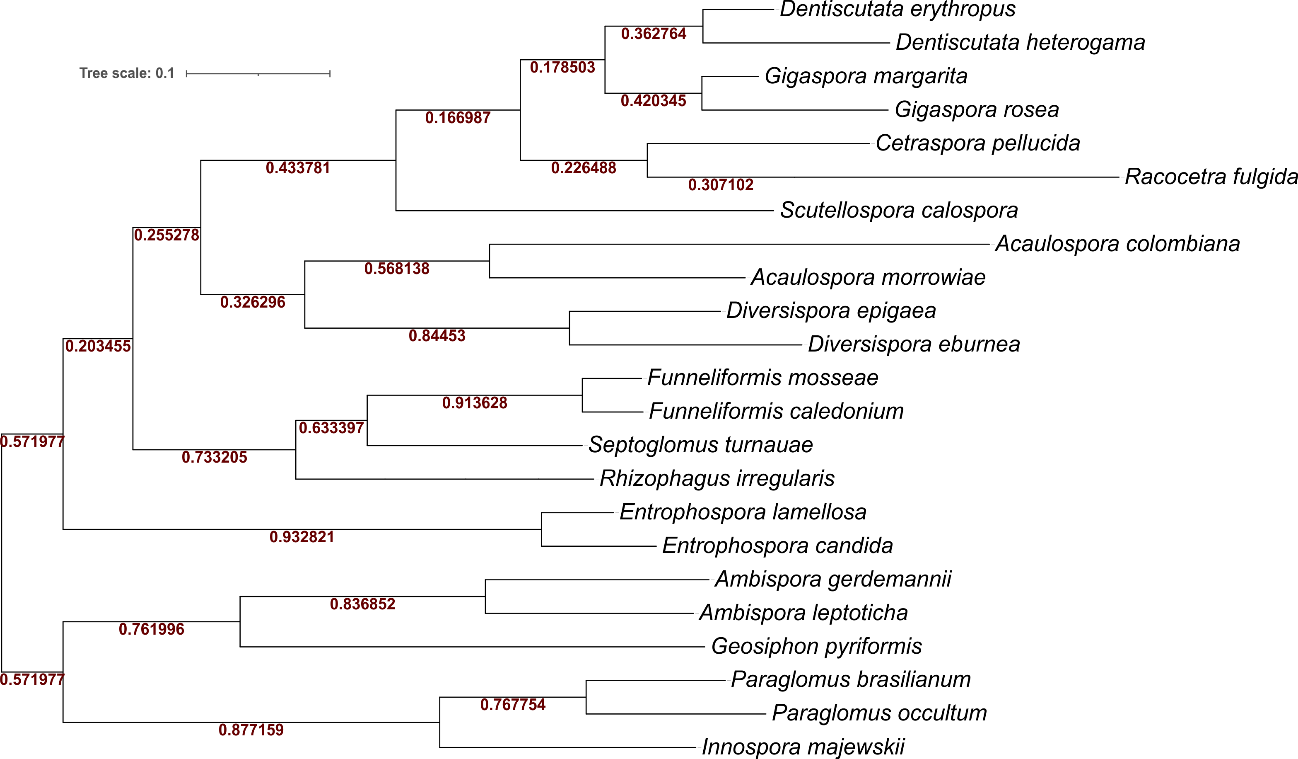

**Figure S1 –** Phylogenetic tree of Glomeromycotina from ortholog genes. The bootstrap support is shown in red. The bar depicts the distance of genetic divergence. The tree agrees with the most recent molecular phylogeny built by Rosling et al., 2024.

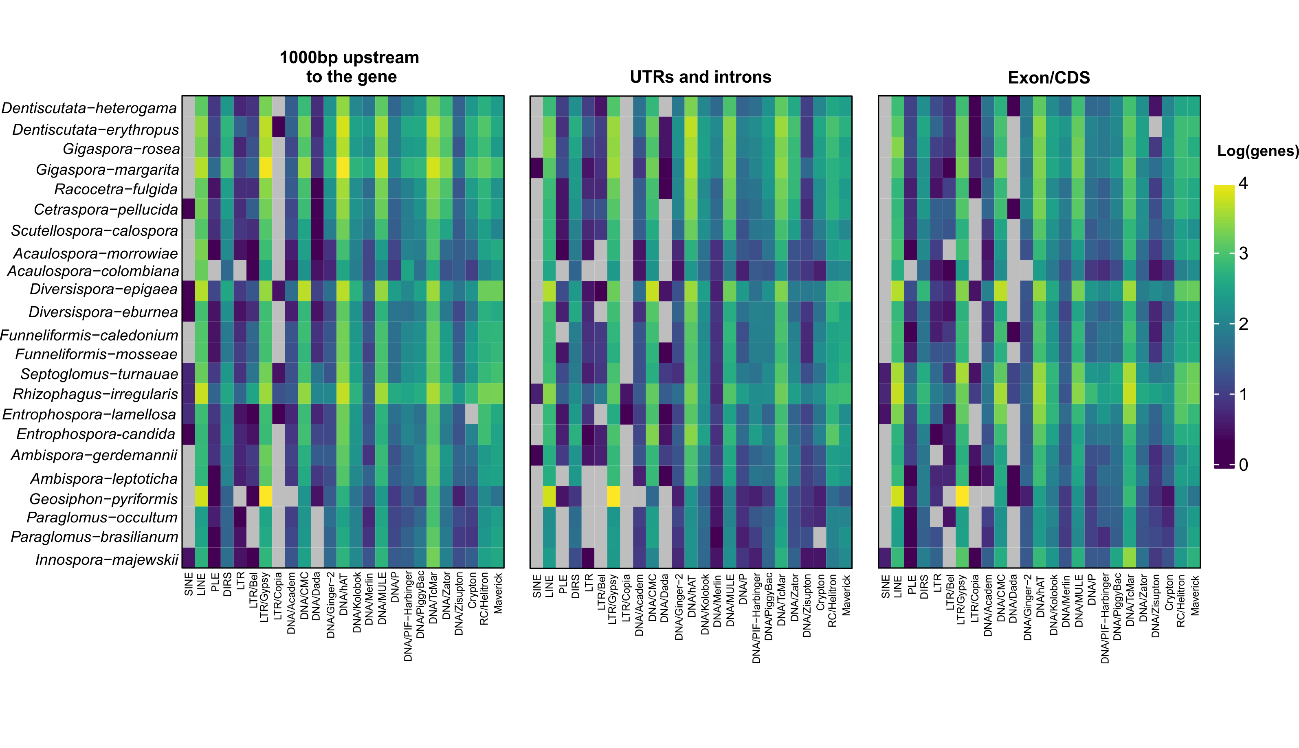

**Figure S2** – Genes associated with TEs in putative promoter region (-1000bp from TSS), in UTRs and introns, and in exons. The gradient of colors represents the number of genes associated with that transposable element in each column. The heatmap are expressed in log scale to minimize the bias of certain species having more genes. In other words, the plots show the relative abundance of genes associated with different TEs in different locations.
